# Molecular effects of indoor tanning

**DOI:** 10.1101/2024.06.04.597225

**Authors:** Pedram Gerami, Bishal Tandukar, Delahny Deivendran, Shantel Olivares, Limin Chen, Jessica Tang, Tuyet Tan, Harsh Sharma, Aravind K Bandari, Noel Cruz-Pacheco, Darwin Chang, Annika Marty, Adam Olshen, Natalia Faraj Murad, Jing Song, Jungwha Lee, Iwei Yeh, A. Hunter Shain

## Abstract

**Background:** Tanning bed users have a significantly increased risk of melanoma, but it remains unclear how indoor tanning drives melanomagenesis. Tanning bed radiation is often thought of as a substitute for natural UV radiation despite differences in the maximum doses, UV content, body sites exposed, and patterns of melanoma that arise.

**Methods:** To better understand the epidemiologic trends and etiology of melanoma associated with tanning bed use, we described the patterns of melanoma in patients with quantifiable tanning bed usage and performed exome sequencing of 182 melanocytes from normal skin of a subset of these patients.

**Results:** Tanning bed users were more likely than non-users to have melanoma on body sites with low cumulative levels of sun damage and were more likely to have multiple melanomas. The melanocytes in normal appearing skin from tanning bed users had higher mutation burdens, a higher proportion of melanocytes with pathogenic mutations, and distinct mutational signatures. These differences were most prominent over body sites that experience comparatively less exposure to natural sunlight.

**Conclusions:** We conclude that tanning bed radiation induces melanoma by increasing the mutation burden of melanocytes and by mutagenizing a broader field of melanocytes than are typically exposed to natural sunlight. The unique signatures of mutations in skin cells of tanning users may be attributable to the distinct spectra of radiation emitted from solariums.

## Introduction

Melanoma is responsible for an estimated 11,000 deaths annually in the United States^1^. The main cause is exposure to ultraviolet radiation, which generates mutations in melanocytes, driving their transformation to melanoma. UV radiation naturally comes from sunlight, but can also be delivered from artificial sources, such as tanning beds. Ever-users of tanning beds are at higher lifetime risk of melanoma^2^, and their melanomas occur at an earlier age^2,3^. The incidence of melanoma has been rising for decades, likely in part from increased screening^4^. However, the rising incidence of melanoma has also coincided with an uptick in tanning bed usage and has disproportionately affected young women, the main clients of the tanning industry. Population-based studies have shown a spiked incidence of melanoma in young women following periods of increased tanning bed usage nationally, suggesting some of the increase in melanoma among this demographic is attributable to indoor tanning^5,6^ Based on the available evidence, the American Academy of Dermatology opposes indoor tanning^7^, and the World Health Organization (WHO) classifies tanning beds as a group 1 human carcinogen, similar to asbestos or cigarette smoke^8^. Despite these warnings, 30 million people, including 2.3 million adolescents, utilize indoor tanning annually in the United States^9^.

Popularity and support of indoor tanning is bolstered by our incomplete understanding of the effects of artificial UV radiation, providing opportunities for the solarium industry to market their product in spite of its link to skin cancer. For instance, the ultraviolet radiation that reaches the earth’s surface is 95% UVA and 5% UVB^10^, while tanning beds typically have even higher proportions of UVA relative to UVB^9^. The tanning bed industry argues that indoor tanning is safer than natural sunlight because UVB is more mutagenic than UVA^11^; however, the spectral irradiance of tanning beds and outdoor sunlight is similar in the UVB range and 10 to 15 times higher in the UVA range than outdoor sunlight, counteracting their argument^12^. Moreover, the tanning industry has marketed the pre-vacation tan as a safe way to photo-adapt skin in anticipation of recreational exposure^13^, supported by observations that the relative risk of melanoma is higher in people who have a history of intermittent, blistering sunburns than in outdoor workers, experiencing daily UV radiation^8^. However, people who burn easily tend to avoid occupations with intense sun exposure, and most outdoor workers wear protective clothing over their trunk and upper extremities, where melanoma is most common.

Considering the high stakes of tanning bed exposure, research is needed to unambiguously illuminate the effects of tanning bed radiation on skin cells, to elucidate how tanning bed usage drives melanomagenesis. Towards this goal, we performed a case/control analysis of patients with/without a history of tanning bed use and a molecular analysis of skin cells from a subset of these patients.

## Results

### Associations between tanning bed usage and melanoma

To better understand the relationship between melanoma and indoor tanning, we interrogated the medical records of 32,315 patients seen by the Dermatology service at Northwestern University (Fig. 1A). Among these patients, 7474 had a self-reported history of tanning bed usage. The extent of tanning bed usage was quantifiable for 2,934 patients, forming our “case” cohort. Among 24,841 patients with no history of tanning bed usage, we randomly selected a subset of 2,929 patients, age-matched to the tanning bed cohort, to serve as the “control” cohort. The tanning cohort was more likely to be female, have a history of sunburn, a history of heavy sun exposure, and a family history of melanoma (Table S1). The incidence of melanoma in our tanning bed cohort was 5.1% compared to 2.1% in the control group (*p*<0.0001, chi-squared test). Multiple logistic regression analysis showed tanning bed use was associated with an increased risk for melanoma (odds ratio of 2.0, 95% confidence interval 1.38, 2.98), after adjusting for age, family history of melanoma, and sunburn history. Further, there was a dose-dependent relationship between number of tanning bed exposures and relative risk for melanoma (*p*<0.0001) (Fig. 1B).

**Figure 1.**
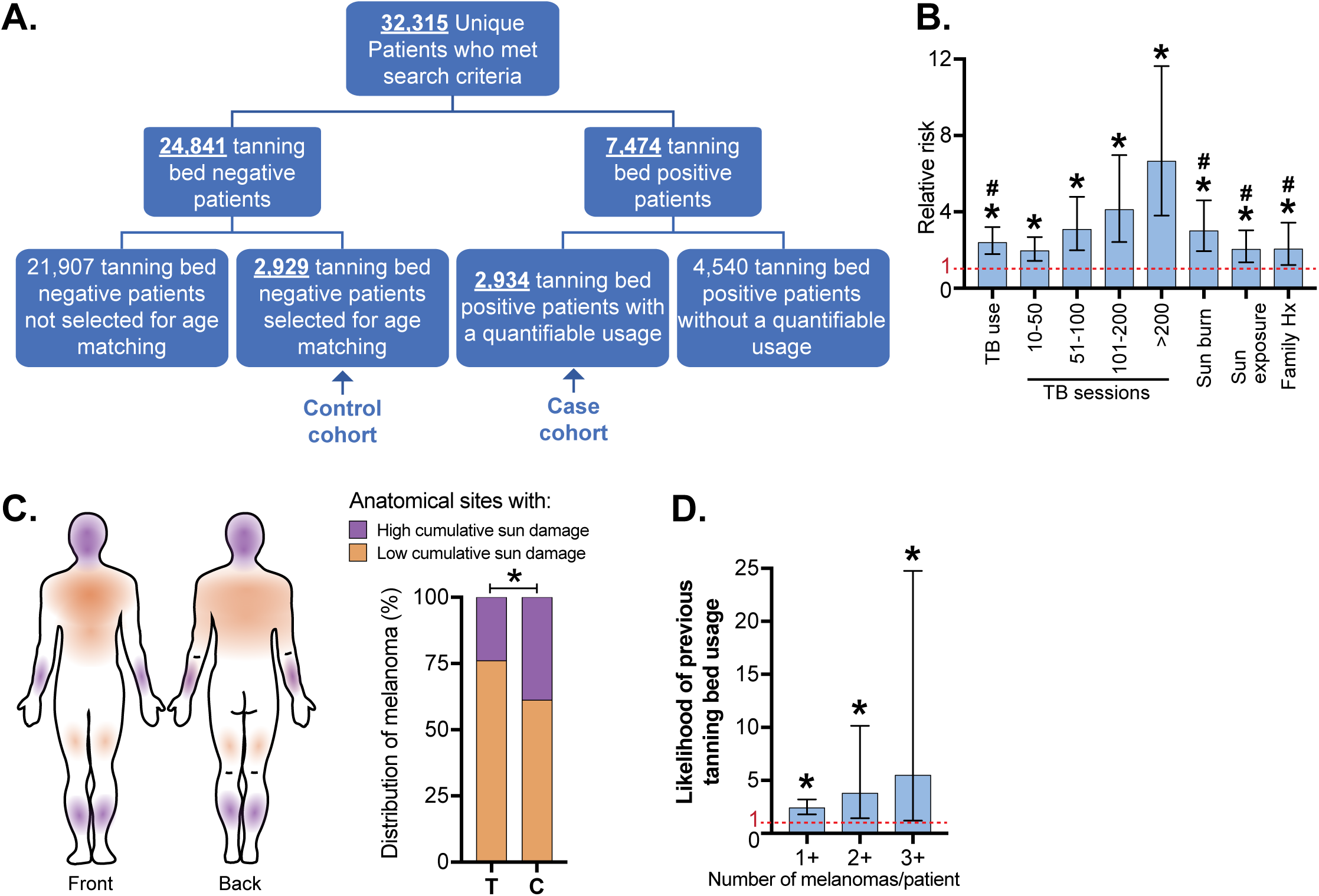
Tanning bed users are more likely than non-users to have multiple melanomas on body sites with low cumulative sun damage. **A.** A case/control cohort was generated from patients seen at a high-risk skin cancer clinic at Northwestern University. Only patients with quantifiable tanning bed use were included in the case cohort, and an equal number of age-matched controls were randomly selected. **B.** The relative risk (RR) of melanoma associated with: tanning bed (TB) usage (including subsetted data based on the number of tanning sessions), history of sun burn, history of heavy sun exposure, and family history of melanoma. Asterisk (*) and number sign (#) denote significant deviance from RR value of 1 (*p* < 0.05), respectively in univariate or multivariate analyses (see methods for details on statistical tests). **C.** The anatomic distribution of melanomas on body sites with low- or high-cumulative sun damage. T and C indicate the tanning and control cohorts. Asterisk indicates *p*-value less than 0.05 (Student’s t-test). **D.** The likelihood of previous tanning bed usage in patients, stratified by the number of melanomas that they have had. Asterisk indicates significant deviance from RR value of 1 (Chi-squared test).

Melanomas in tanning bed patients had a different anatomic distribution than melanomas from the control cohort. The World Health Organization (WHO) distinguishes melanomas that arise on skin with high cumulative sun damage versus melanomas on skin with low cumulative sun damage^14^. Melanoma was more common on body-sites with low cumulative sun damage in tanning bed users compared to non-users (Fig. 1C). A previous study observed a similar anatomic distribution of melanoma in tanning bed users, though their case-control cohort was not matched for age^15^. Multiple primary melanomas were also more common in tanners than non-tanners (Fig. 1D). Based on these observations, we hypothesized that indoor tanning increases the risk of melanoma in two ways: first, by increasing the mutation burden of melanocytes, and second, by mutagenizing a larger field of melanocytes, beyond the body sites that are typically exposed to natural sunlight, creating a broader field effect. To test this hypothesis, we compared the mutational landscapes of melanocytes from normal skin samples of tanning bed users to non-users.

### Molecular consequences of tanning bed usage

Shave biopsies of normal skin were collected from the lower backs or upper backs of 11 tanning bed users. Tanning bed users filled out a questionnaire, modeled after the United Kingdom (UK) Biobank, to assess past histories of sun exposure and other risk factors for skin cancer^16^. They had self-reported histories of extreme tanning bed usage – ranging from 50 to over 750 lifetime sessions, among other risk factors summarized in Table S2.

Normal skin samples were also collected from two different control groups (Fig. 2A) for comparison. In the first control group, 9 patients were recruited from the same high-risk skin cancer clinic as the tanning cohort and matched for age, sex, and risk profiles of the tanning cohort, based on responses to the UK Biobank questionnaire. The tanning cohort was more likely to have been diagnosed with melanoma and to have a history of tanning bed usage, but no other risk factors for skin cancer were significantly different between the groups (Table S2, Fig. S1A). While the tanning cohort and first control group were matched for common risk factors, when they were compared to the participants in the UK Biobank, both were more likely to have a history of melanoma, red or blonde hair, a history of sunburn, and poor tanning ability (Fig. S1B), likely because they were recruited from the same high-risk skin cancer clinic.

**Figure 2.**
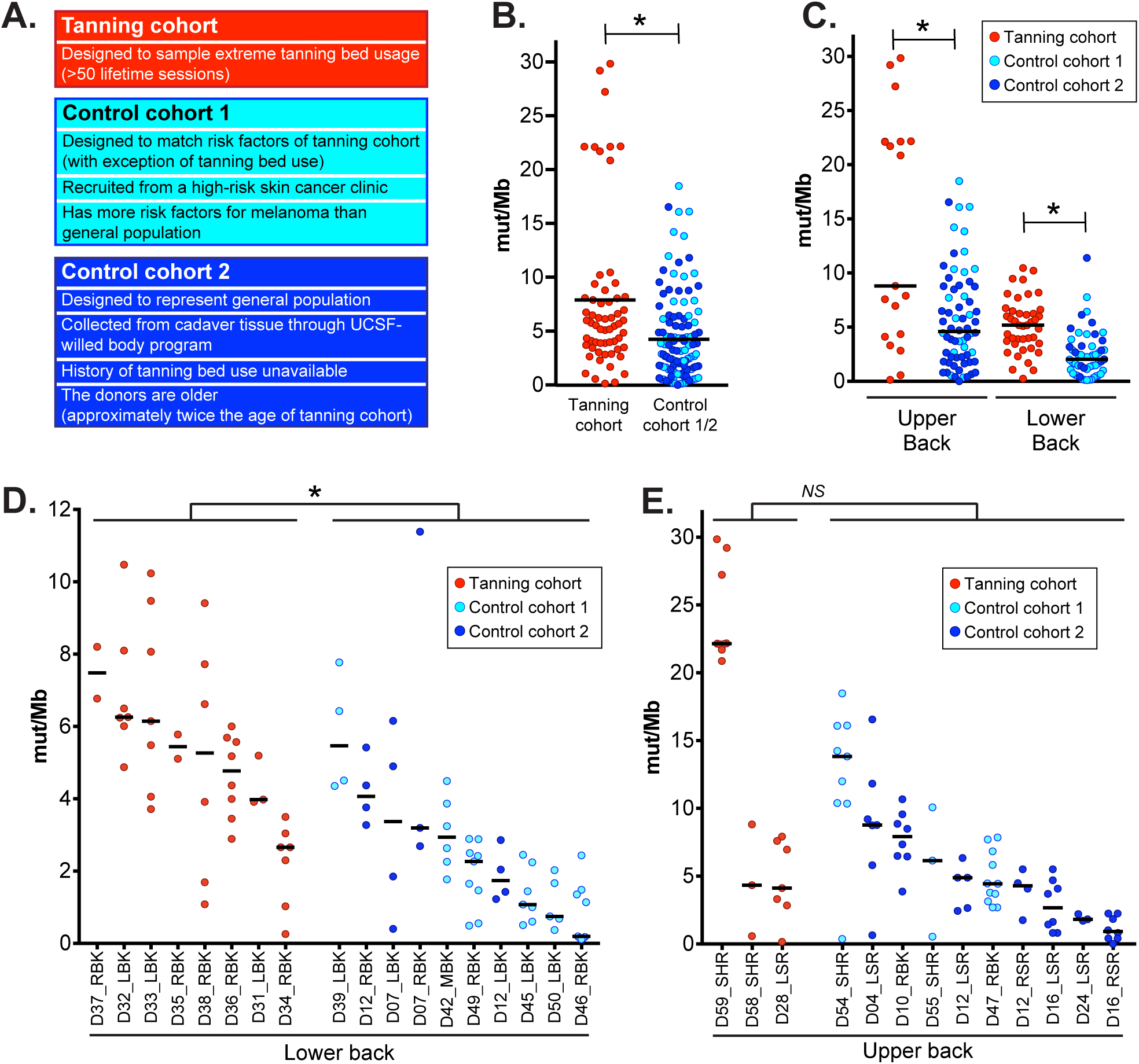
High mutation burdens in melanocytes from tanning bed users. **A.** An overview of the tanning and control cohorts. In panels **B**-**E**, each data point corresponds to the mutation burden (measured in mutations per megabase) of an individual melanocyte with black bars indicating median mutation burdens. Asterisks denote *p-*values less than 0.05 (Wilcoxon rank-sum test). **B.** A comparison of all melanocytes from tanning bed donors to all melanocytes from control donors. **C.** A comparison of melanocytes from tanning bed donors to control donors, separately for each anatomic site. **D.** A comparison of biopsy mutation burdens from tanning bed donors to control donors on the lower back. The mutation burden of each biopsy was calculated from the median mutation burden of its constituent melanocytes. **E.** A comparison of biopsy mutation burdens from tanning bed donors to control donors on the upper back.

For a second control group, we collected normal skin from six cadavers through the UCSF-Willed Body program (Table S2, Fig. 2A). We assumed the cadaver tissue, which included two donors of dark skin tone, would be more representative of the general population than the donors recruited from a high-risk skin cancer clinic. A limitation to this control group is that donors were nearly twice the age of the tanning cohort (78.3 versus 43.6 years of age on average). As another limitation, we were unable to interrogate the past histories of tanning bed usage from the deceased donors, though extreme tanning bed usage (>50 sessions) is rare in the general population.

After collecting normal skin samples, we measured somatic mutations at single-cell resolution from a total of 182 melanocytes derived across these donors (Table S3). It is challenging to comprehensively and accurately detect mutations in an individual cell. Therefore, we developed a protocol to overcome this obstacle, as previously described^17^. We established melanocytes in tissue culture from epidermal biopsies and clonally expanded individual melanocytes to form small colonies, with a median of 213 cells per colony. DNA and RNA was extracted from each colony and further amplified *in vitro*. Exome and transcriptome sequencing was performed on the amplified DNA/RNA from each colony. We developed a bioinformatic workflow to root out amplification artifacts, permitting the detection of somatic mutations at high specificity. Mutation calls were internally benchmarked for each cell, in part, by assessing agreement in mutation support from DNA and RNA of the same cell within highly expressed genes, showing an average detection of 95.2% sensitivity and 97.1% specificity (Table S3).

The mutation burden of melanocytes in tanning bed users was significantly higher than the mutation burden of melanocytes from control donors (Fig. 2B). This trend remained significant when we separately compared melanocytes by anatomic site (Fig. 2C). The mutation burdens of cells from both control groups were less than the cells in the tanning cohort (Fig. S2A). Finally, the tanning cohort had a significantly higher mutation burden at the cell level in a mixed-effect model that adjusted for anatomic site and included a random effect for subject (*p* = 0.0128, see methods).

We also compared mutation burdens at the biopsy level. We defined the mutation burden of a biopsy as the median mutation burden of its constituent melanocytes. Our statistical power was more limited in these comparisons because there were fewer independent biopsies than individual cells. On the lower back, tanning bed biopsies had significantly higher mutation burdens than control biopsies (Fig. 2D, S2B). The difference was not statistically significant on the upper back (Fig. 2E, S2C), though cell-level differences were significant at this site (Fig. 2C). The upper back is the most common site of sunburn^18^ and melanoma^19^, underscoring the vulnerability of this site to sun damage. Tanning-bed-induced mutations would be expected to contribute to a smaller proportion of mutations in the underlying skin cells from the upper back, likely requiring a larger sample size to detect a statistically significant difference at the biopsy level.

Given that tanning bed users experience higher doses and different blends of UV radiation than typically encountered from natural sunlight, we performed mutational signature analyses^20^ to interrogate the types of mutations in their melanocytes. The dominant mutational signature in cells from both tanning bed users and non-users was signature 7. Signature 7 is characterized by cytosine to thymine (C>T) mutations with a pyrimidine upstream of the mutant basepair (Fig. S3A), and it has been attributed to UV-radiation-induced damage^20^. Cells from tanning bed users had higher proportions of mutations associated with signature 7, but the difference was not significant (Fig. 3B).

**Figure 3.**
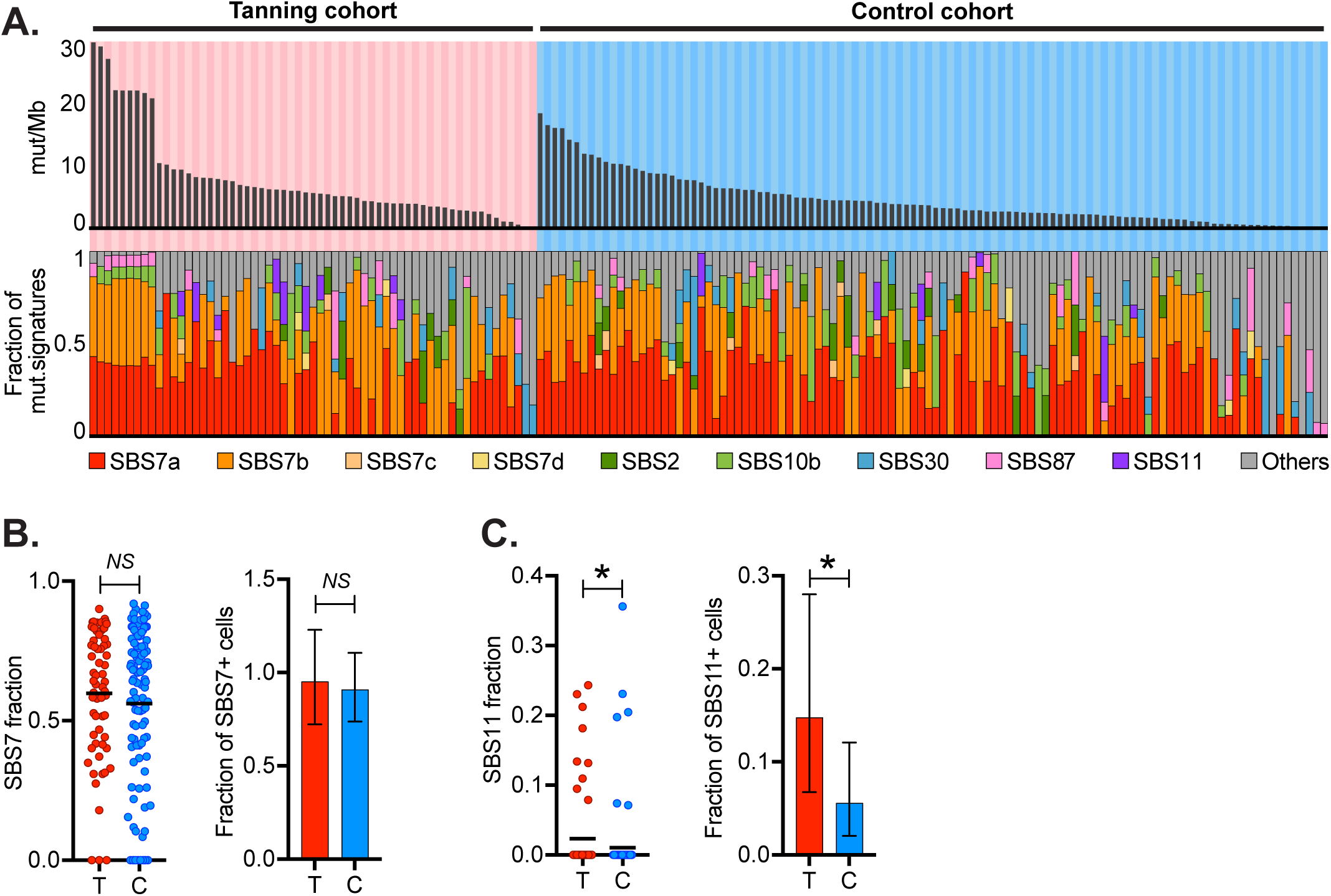
Distinct types of mutations between melanocytes in tanning bed donors versus control donors. **A.** The top panel depicts the mutation burden (mut/Mb) of melanocytes with each column representing a single melanocyte, arranged in descending order for each cohort. The middle bar graph shows the fraction of mutations from UV radiation (CC>TT or (C/T)C>T). The bottom panel shows the fractions of different mutation signatures for each melanocyte. **B.** The left scatter plot shows proportions of signature 7 in melanocytes from the tanning cohort (T) and control cohort (C) (*p*-value: 0.609, Wilcoxon rank-sum test). The right bar graph compares the fraction of melanocytes with any detectable signature 7 in each cohort (*p*-value: 0.6867, Poisson test). **C.** Data is plotted as described in the previous panel but for signature 11 (*p*-value: 0.0405, Wilcoxon rank-sum test on the left, and *p*-value: 0.00813, Poisson test on the right).

Signature 11 was the only signature to reach a statistically significant difference between the cohorts (Fig. 3C). Signature 11 shows similarities to signature 7 in that it is characterized by C>T mutations, but in contrast to signature 7, there is a pyrimidine downstream of the mutant basepair (rather than upstream, Fig. S3B). In the current catalogue of mutational signatures (COSMIC v. 3.4, Fig. S3D), melanoma is the most common tumor subtype associated with this signature. The tanning history of melanomas included in the catalogue of mutational signatures is unknown, but it is likely that a subset arose from patients with a history of indoor tanning. Signature 11 was originally attributed to temozolomide treatment, based on an anecdotal association with glioblastoma pretreated with temozolomide^21^, however, *in vitro* studies have found that temozolomide induces a different mutational profile^22^. Currently, the etiology of signature 11 is unknown. Considering that signature 11 is most common in UV-radiation-induced cancers, enriched here in skin cells of tanning bed users, and it shows similarities to canonical UV-radiation-induced mutations, it may be attributable to unique blends of UV wavelengths experienced by indoor tanning users, but further studies are warranted.

Melanocytes from physiologically normal skin can harbor mutations known to drive melanoma. These cells are potential precursors to melanoma but require additional genetic alterations and/or microenvironmental stimuli to grow out and form clinically detectable neoplasms. We found 40 pathogenic mutations in 23 unique melanocytes (Fig. 4A). Most driver mutations, observed here, were predicted to activate the MAPK signaling pathway, in which loss-of-function mutations affecting the *NF1* tumor suppressor gene were especially common. Melanocytes from tanning bed users were more likely to have a pathogenic mutation than melanocytes from control donors, and melanocytes from the upper back were more likely to have a pathogenic mutation than those from the lower back (Fig. 4B-C).

**Figure 4.**
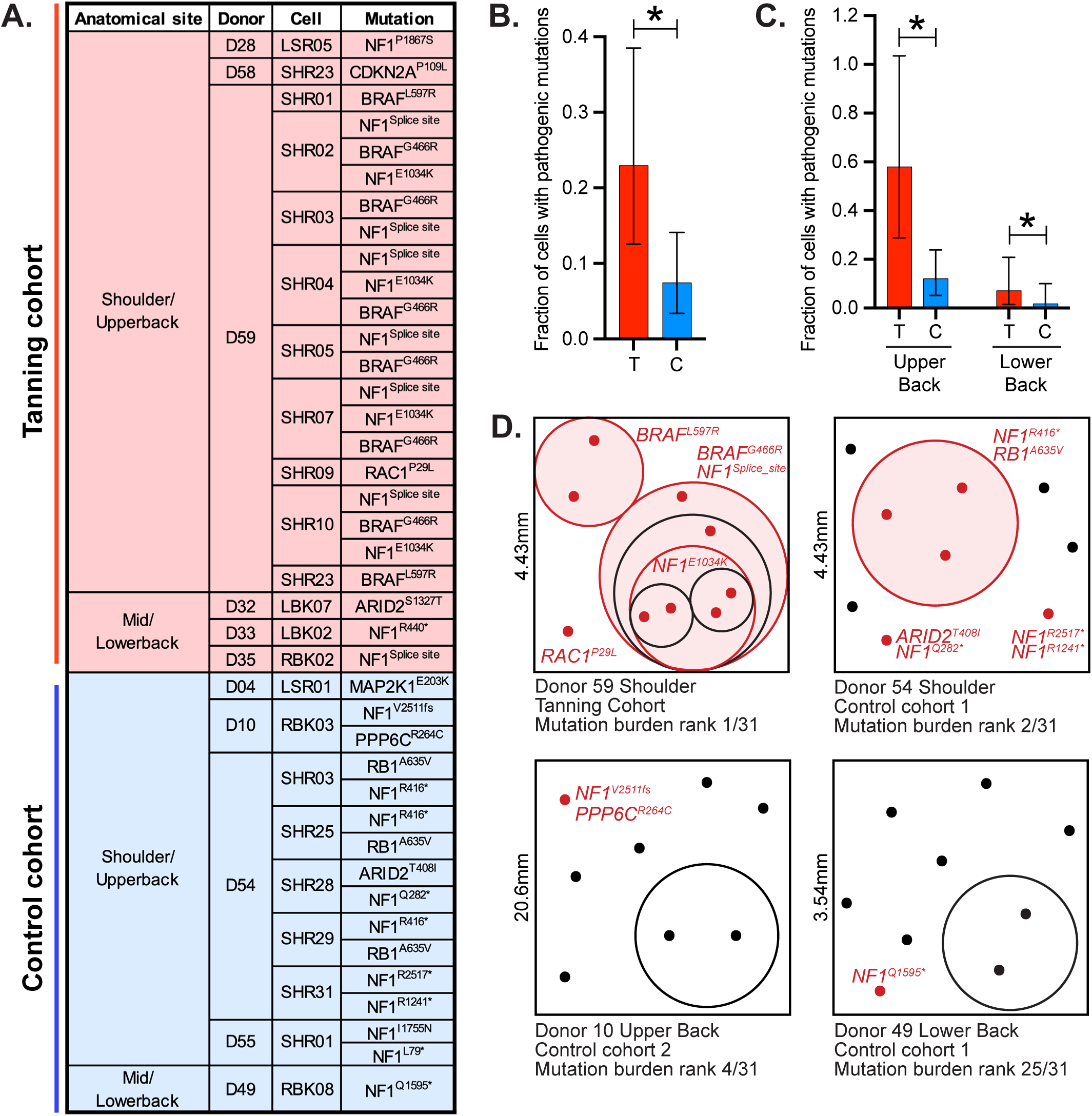
Pathogenic mutations are common in melanocytes from tanning bed users. **A.** A list of pathogenic mutations observed, broken down by cohort, body site, donor, and cell. **B.** The fraction of cells with pathogenic mutations in tanning bed users (T) or control donors (C). Asterisk indicates *p*-value less than 0.05 (Poisson test) with error bars showing 95% confidence intervals. **C.** Data is plotted as in panel B, but here, cells are broken down by anatomic site. **D.** Four biopsies had 2 or more melanocytes with shared subsets of mutations. In these schematics, each dot represents an individual melanocyte, and phylogenetically related melanocytes are circled. Melanocytes with pathogenic mutations are noted in red with their driver mutations labeled. The true spatial localization of each cell, within biopsies, is unknown, but these schematics attempt to illustrate field sizes and clonal distribution of skin biopsies.

Some melanocytes shared subsets of mutations, indicating a phylogenetic relationship (Fig. 4D). These melanocytes likely descended from fields of clonally related cells present in the skin. We were underpowered to directly establish a relationship between tanning bed usage and clonal structure, however, biopsies with high mutation burdens were more likely to have a field of melanocytes (Fig. 4D). We propose that UV radiation (whether it comes from natural or artificial sources) produces fields of melanocytes within human skin in two ways. First, UV radiation introduces pathogenic mutations into a subset of melanocytes, allowing these cells to outcompete neighboring wild-type cells. Second, UV radiation introduces deleterious mutations into some melanocytes, allowing neighboring cells to passively expand in their stead.

## Discussion

Our study was motivated by a clinical presentation recurrently seen in Dermatology clinics where tanning bed usage is especially high. Young tanning bed users, without a family history of melanoma, periodically develop multiple melanomas on body sites that receive low cumulative sun damage. The presentation of tanning-bed induced melanomas is reminiscent of familial melanoma^23^. In families with a strong predisposition to melanoma, each person inherits one “hit” in their germline DNA (e.g. a *CDKN2A* mutation) bringing all their melanocytes one step closer to melanoma. Like tanning bed users, patients with familial melanoma also develop disease at a young age, are more likely to get multiple melanomas, and a larger field of melanocytes are at risk for transformation^23^. We hypothesized that tanning beds mimic these circumstances by mutagenizing a large surface area of the body, beyond the sites typically exposed to natural sunlight. Indeed, body sites that typically receive low cumulative sun damage are intensely exposed during tanning sessions. This constitutes nearly 1.5X the body surface area of heavily sun-exposed skin (Fig. S5)^24,25^, significantly increasing the skin surface at high risk of melanoma in tanning bed users (Fig. 5).

**Figure 5.**
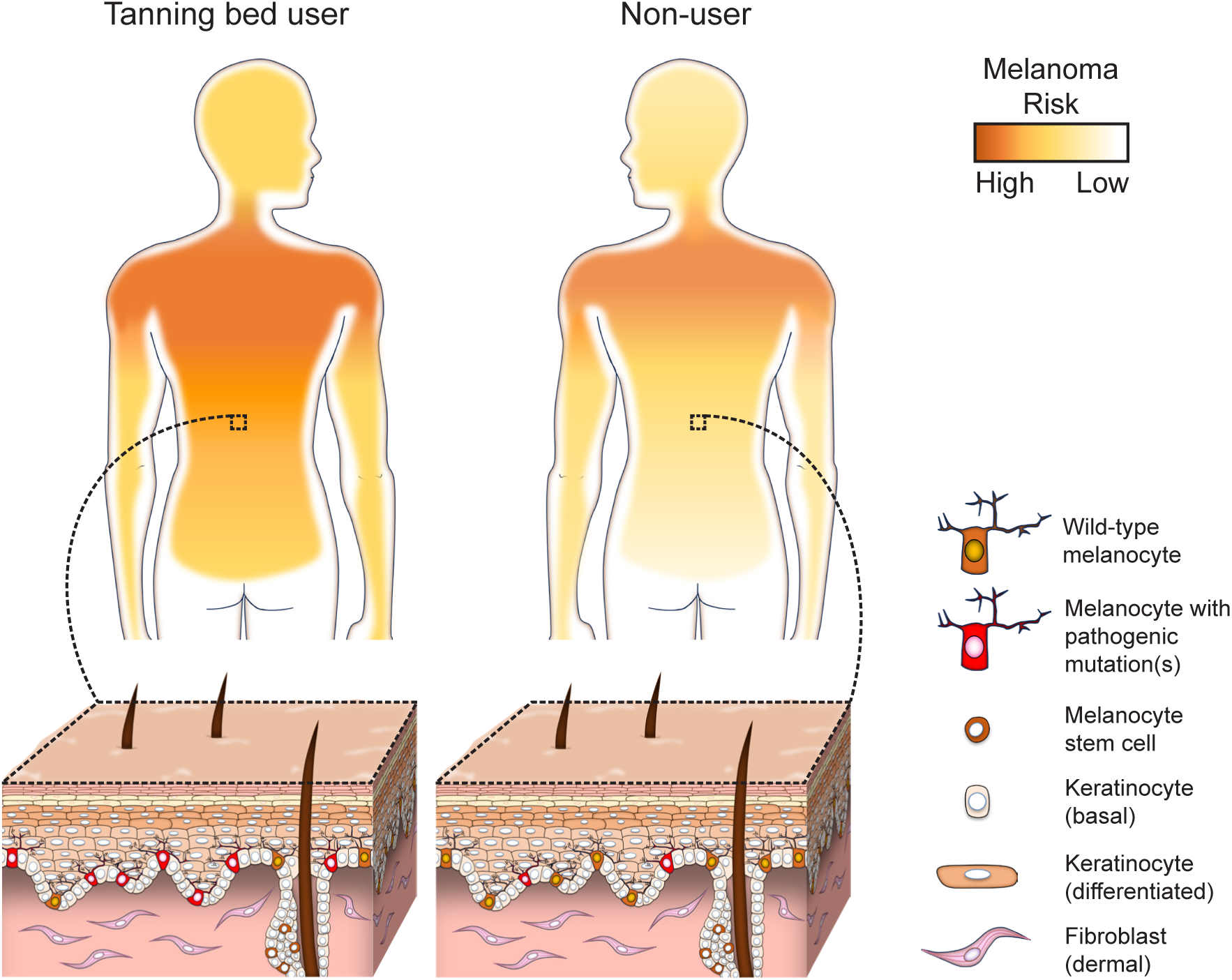
High proportion of melanocytes in tanning bed users have pathogenic mutations, increasing risk of melanoma, particularly over body sites with low cumulative sun damage.

Young patients, with an extreme history of tanning bed usage, had more mutations in their cutaneous melanocytes than donors who were nearly twice their age from the UCSF Willed-Body program. Tanning bed users also had more mutations than demographically matched donors at similarly high risk of skin cancer. The difference in mutation burdens was most glaring on the lower back – a body site that receives comparatively less damage from natural sunlight but is exposed in tanning beds. In addition to having higher numbers of mutations, melanocytes from tanning bed users were more likely to have a pathogenic mutation than melanocytes from either control group (Fig. 5). Having a high fraction of melanocytes with pathogenic mutations would be expected to increase one’s risk of melanoma and potentially multiple primary melanomas by generating a large pool of precursor cells one step closer to transformation, akin to patients with heritable forms of melanoma. In our experience, among patients with multiple primary melanomas, tanning bed use is a more common etiologic factor than high penetrant germline mutations^26^.

There were subtle differences in the types of mutations in melanocytes between tanning bed users and control donors. Tanning bed users were more likely to have a pyrimidine immediately downstream of the mutant site, indicated by a higher proportion of cells with signature 11 mutations. These differences may result from the unique blends of radiation experienced by tanning bed users, with much higher levels of UV-A radiation than natural sunlight. However, more work is needed to understand the source of these mutations. One signature was notably absent. UV-A radiation can induce a reactive oxygen species (ROS) signature^27^, but these signatures were not observed here. It is possible that ROS-induced mutations are difficult to detect amid the large numbers of UV-radiation-induced mutations. Alternatively, physiological levels of ROS may not reach mutagenic thresholds *in vivo*.

Our work provides new insights to inform public health guidance on indoor tanning. Given the high levels of mutational damage in skin cells from tanning bed users, it is difficult to justify marketing claims that the spectra of UV radiation in solariums are safer than natural sunlight. Another popular claim by tanning advocates is that a pre-vacation tan can photo-adapt skin in anticipation of recreational sun exposure. However, we see that tanning bed usage raises the mutation burden and risk of melanoma particularly in skin cells that receive low cumulative sun damage. In closing, tanning bed exposures are often thought of as a substitute for natural UV radiation despite differences in the maximum doses, UV content, body sites exposed, and patterns of melanoma that arise. Our work highlights unique ways in which tanning beds shape the mutational landscapes of skin cells, helping to explain the distinctive patterns of melanoma in this patient population.

## Acknowledgements

The study was supported by grants from: NIH NCI (R01 CA265786), NIH NIAMS (AR080626), Department of Defense Melanoma Research Program (ME210014), Melanoma Research Alliance (Team Science Award and Dermatology Fellows Award), the LEO Foundation Region Americas Award, Cancer Center Support Grant P30CA082103, the IDP Foundation, and the Greg and Anna Brown Family Foundation.

---

**Figure S1. Control cohort 1 donors match the risk profiles of tanning bed donors, however, both groups are at higher risk of skin cancer than the general population.** Donors were recruited for molecular studies of their skin cells and asked to fill out questions, modeled after the UK Biobank, surveying their past exposure to UV radiation and other risk factors for skin cancer. See Table S1 for the full wording of each question and answer. Bar graphs summarize the responses to each question with the risk trait colored more darkly. Asterisk denotes *p*-values less than 0.05 (Chi-squared test). Panel **A** compares the responses of the tanning cohort to the control cohort. Panel **B** compares the responses of both the tanning/control cohorts to the UK Biobank participants, chosen to represent the general population.

**Figure S2. Comparison of mutation burdens of melanocytes from tanning bed users to each control cohort separately.** Each data point corresponds to the mutation burden (measured in mutations per megabase) of an individual melanocyte. Black bars indicate median mutation burdens. Asterisks denote *p*-values less than 0.05 and *NS* indicate not significant (Wilcoxon rank-sum test). **A.** A comparison of melanocytes from tanning bed donors to melanocytes from each control cohort, separately for each anatomic site. **B.** A comparison of biopsy mutation burdens from tanning bed donors to control donors from each cohort on the lower back. The mutation burden of each biopsy was calculated from the median mutation burden of its constituent melanocytes. **C.** A comparison of biopsy mutation burdens from tanning bed donors to control donors from each cohort on the upper back.

**Figure S3. Composition of mutational signatures.** 96 bar plots show the fraction of different mutations (y-axis) in all possible tri-nucleotide contexts (x-axis) for signature 7 (panel **A**) and signature 11 (panel **B**). Panel **C** compares the trinucleotide contexts of the tanning cohort (top panel), control cohort 1/2 (middle panel), and the difference between these two cohorts (bottom panel). Panel **D** illustrates the mutational burden attributable to signature 11 in the current catalog of COSMIC signatures (version 3.4).

**Figure S4. Phylogenies of melanocytes from four skin biopsies with clonally related cells.** Phylogenetic trees were constructed for individual melanocytes from four skin biopsies. Each tree is rooted in the germline state. Trunk and branch distances are proportional to the number of shared and unshared somatic mutations between cells. The identity of each cell is labeled at the terminus of each tree, and pathogenic mutations are labeled in red. Some cells were sequenced with exome baits, and some were sequenced with a smaller gene panel, as indicated. See figure 4D for a series of schematics portraying the potential clonal structure of each biopsy.

**Figure S5. Total body surface area in the regions with high (purple) and low (orange) cumulative sun damage (CSD).** The figure is based on Lund and Browder chart. The high CSD regions cover 35% and low CSD regions cover 52% of the body area. The remaining anatomical regions are considered sun shielded.

## Methods

### Epidemiological Analyses (related to figure 1 and table S1)

Under Northwestern University Institutional Review Board protocol STU00211546, patient records collected from the Department of Dermatology at Northwestern Medicine were evaluated using an EDW (electronic data warehouse) consolidated database based on the following inclusion criteria: aged 18-70 before 2019, a quantifiable history of tanning bed use, at least 1 dermatology visit before 2019, and at least 1 additional visit 3 or more years prior to or after the first visit. A total of 2934 patients met these criteria and were sorted into the positive tanning bed cohort. The control cohort included a random selection of 2934 age matched patients with no prior history of tanning bed use. Of the 2934 control patients, 5 were excluded due to duplicate patient records. Both family and personal history of melanoma, as well as sunburn and heavy sun exposure history were collected for each cohort. For the multiple logistic regression analysis, the final cohort included 2932 tanning bed positive patients and 2926 tanning bed negative patients. Heavy sun exposure was excluded as a factor of the multiple logistic regression due to several patients missing this data point.

Statistical analyses were performed in Microsoft Excel with XLSTAT add-in to perform chi-square test to compare associations in categorical variables and Student’s t-test was used to compare mean values. All tests were two sided. In addition, relative risk was also calculated for all included risk factors and significance was calculated using a z score. A *p*-value of <0.05 was considered statistically significant. A multiple logistic regression was performed using SAS version 9.4.

### Skin biopsy collection for molecular analyses

Donors were recruited from the Northwestern Dermatology high-risk skin cancer clinic for the tanning cohort and first control cohort (see figure 2A for an overview of all sample cohorts used in molecular analyses). For the tanning cohort, eleven donors were recruited, and each had greater than 50 self-reported tanning sessions. No other inclusion criteria were put in place, and since tanning usage is most common in young women, the cohort skewed towards this demographic. We also asked the donors in the tanning cohort to fill out a questionnaire related to their previous exposures to UV radiation and risk factors for skin cancer. To generate the questionnaire, we identified the subset of questions from the UK Biobank that related to skin cancer risk and asked our donors to answer them as well. This strategy allowed us to put their responses in relation to the population in the UK Biobank. Donors in control cohort 1 were recruited from the same patient pool at Northwestern and selected to match the sex, age, and risk profiles of the tanning cohort with the exception of tanning bed usage. All donors in control cohort 1 self-reported no tanning bed usage. Both the tanning and control cohorts underwent additional scrutiny to contextualize them within the general population. This was achieved by comparing their responses to risk factor questionnaires with publicly available responses from participants of the UK Biobank survey (Fig. S1). Informed consent was obtained from all donors recruited at Northwestern. This study was approved by the Institutional Review Board protocol STU00009443.

Donors from control cohort 2 were collected from the UCSF Willed Body program. The UCSF Willed Body program was established to receive the remains of individuals who choose to donate their body for medical research. All donors consented, as part of their living will, prior to their death. There were no inclusion criteria for Willed Body donors, though most donors through this program are of advanced age. All biopsies analyzed in this study were obtained from the upper back (including shoulders) and lower back (including mid-back) regions.

### Skin biopsy preparation for single-cell genotyping

Shave biopsies, ranging from 3-5mm in their longest dimension, were collected from living donors or cadaver tissue, placed in saline, put on ice, and transported to the Shain laboratory at UCSF for molecular analyses. Upon receipt, skin biopsies were treated with dispase (10mg/ml) for 16 hours, breaking up collagens connecting the dermis to the epidermis. After incubation, the epidermis was physically peeled from the dermis, minced with a scalpel, trypsinized (0.05% trypsin for 3 min), and vortexed (every 5-10 seconds for 3 minutes) to establish a suspension of single cells. Suspended cells were placed in tissue culture and grown in CNT40 media (CELLnTEC) + 5% antibiotic-antimycotic. After one week the cultures contain a mixture of melanocytes and keratinocytes. To separate these cell types, trypsinization (0.05%) was performed for 3 minutes, detaching melanocytes while leaving keratinocytes adherent to the plate.

After achieving stability in bulk cell culture, individual cells were seeded into 96-well plates using serial dilution. These plates underwent immediate screening to eliminate wells containing multiple or no cells. The single-cell cultures were maintained until they ceased expanding, typically resulting in colony sizes of 200 cells. Colonies were harvested in RLT buffer (Qiagen, 79216).

We further amplified the genomic DNA and mRNA from each harvested colony using the G&T sequencing protocol^28,29^. The G&T-seq protocol describes how to separate genomic DNA and mRNA. Once separated, the genomic DNA was amplified in vitro via Multiple Displacement Amplification (MDA) (Qiagen REPLI-g Single Cell Kit, 150345) or Primary Template Amplification (PTA) (BioSkryb ResolveDNA Whole Genome Amplification Kit, 100136). We switched to PTA when the technology became available because it has higher fidelity amplification than MDA^30^. The mRNA was amplified with SMART-Seq2 (Switch Mechanism at the 5’ End of RNA Templates 2)^31^. Collectively, the clonal expansion and *in vitro* amplification of nucleic acids produced sufficient genomic material to call somatic mutations from individual melanocytes.

A reference source of normal DNA was collected from each donor. For the living donors recruited at Northwestern, a buccal swab was collected. For the cadaver tissue at UCSF, we biopsied a distant skin sample. DNA from buccal swabs was isolated with prepIT.L2P (DNA Genotek, PT-L2P-5), whereas DNA from skin samples was isolated using the DNeasy Blood & Tissue Kit (Qiagen, 69504).

### Sequencing and somatic alteration calls

Nucleic acids were fragmented to a target size of 350bp using Covaris LE220, followed by end-repair and ligation to IDT 8 or 10 dual index adaptors. Amplification was carried out using the KAPA HyperPrep Kit (KK8504). Subsequently, the genomic DNA libraries were enriched for the exome through hybridization with either NimbleGen SeqCap EZ Exome + UTR (06740294001), KAPA HyperExome v1 (09062556001), or KAPA HyperExome v2 (9718648001) probes using KAPA HyperCapture Reagent kit (09075828001). Paired-end sequencing (either 100 or 150bp) was conducted on Illumina HiSeq 2500 or NovaSeq 6000 instruments.

Sequencing data from genomic DNA was aligned to the hg19 version of the genome with BWA (v2.0.5)^32^ and deduplicated with Picard (v2.1.1). Subsequently, the reads underwent additional curation to realign indels and recalibrate base quality using GATK (v4.1.2.0). For RNA sequencing data, alignment to both the genome and transcriptome was conducted with STAR align (v2.1.0)^33^. The reads were then deduplicated using Picard (v2.1.1) and gene-level read counts were quantified using RSEM (v1.2.0)^34^.

Copy number alterations were inferred from DNA- and RNA-sequencing data using CNVkit (v.0.9.6.2)^35,36^. A candidate set of germline heterozygous SNPs was called with FreeBayes (v.1.3.1)^37^ and further filtered to only include SNPs observed in the 1000 Genome Project and between 40-60% allelic frequency mapping to each allele. A candidate set of short insertions and deletions was called with Pindel^38^. Candidates were filtered to remove those with fewer than 4 supporting reads and which were present in the reference bam. Remaining candidates were manually screened to remove likely alignment artifacts. Somatic point mutations were called as previously described^17^. Briefly, a candidate list of point mutations was called with MuTect by comparing the bam file of each clonal expansion to the reference bam file from the same patient. The MuTect calls included both somatic point mutations and amplification artifacts that arose during MDA or PTA. To remove amplification artifacts, we leveraged patterns in the sequencing data. We searched for supporting reads in the RNA-sequencing data from each clonal expansion, allowing us to validate or invalidate candidates in highly expressed genes. We also interrogated phasing patterns for variants near germline heterozygous SNPs. True mutations occur in complete linkage with at least one parental allele, whereas artifacts tend not to show this pattern. For the remaining variants, which were not in highly expressed genes or near germline heterozygous SNPs, we inferred the likelihood that they were a true somatic mutation based on their allele frequency.

### Mutation burden and mutational signature analyses (related to figures 2, 3, S2, and S3)

Mutation burdens were calculated as mutations per megabase. We counted the number of mutations as described above. To determine the footprint of genome with sufficient coverage in each clonal expansion, we ran the footprints software^17^ to count the precise number of basepairs with 10X coverage or greater. For the analysis of mutational signatures, we compiled somatic mutations across all cells from both cohorts (Table S4) and established trinucleotide contexts for single base substitutions using the Bioconductor library BS.genome.Hsapiens.UCSC.hg19 (v1.4.3). This analysis was restricted to cells harboring a minimum of 10 mutations, as profiles with fewer mutations are statistically unreliable. DeconstructSigs R package (v1.9.0)^39^ was employed to generate the mutation signature profile for each cell using pre-defined 78 COSMIC (v3.4) signatures. These signatures were previously delineated by SigProfiler^40^ and listed in Catalogue of Somatic Mutations In Cancer (COSMIC) database (https://cancer.sanger.ac.uk/signatures/sbs/). A stacked barplot was constructed to illustrate the prevalence of signatures 7a, 7b, 7c, 7d, and the top five additional signatures for each individual cell (Fig. 3A). Remaining signatures were grouped under an "others" category. Specifically, we also compared signatures 7 and 11 between the tanning bed and control cohorts, examining the fraction of these signatures per cell. We employed the Wilcoxon rank-sum test to assess the significance of any observed differences. We further scrutinized signatures 7 and 11 by calculating the fraction of cells positive for each signature in both cohorts. Confidence intervals and *p*-values were computed using the Poisson test.

### Annotation of pathogenic mutations (related to figures 4A-C)

Somatic mutations were manually scrutinized by the authors to annotate mutations known to be under selection in melanoma. The full list of mutations is available in (Table S4), and our pathogenic annotations are shown in column AA of that table. The final list of pathogenic mutations is available in Fig. 4A.

### Clonality analyses (related to figures 4D and S4)

To visually depict the phylogenetic relationship among melanocytes sharing somatic mutations, we created clonality plots, as shown in Fig. 4D. In these plots, the size of each square corresponds to the surface area of the skin biopsy at the indicated scales. To simplify the visualization, the schematics depict skin biopsies as squares, even though their shapes varied. For punch or shave biopsies, the area was calculated based on their known diameters. In cases of larger biopsies with irregular shapes, images were captured with a scale and analyzed using ImageJ^41^ to determine the area. Melanocytes were represented as randomly positioned dots, with clonally related ones enclosed within circles.

The total surface area represented by each melanocyte was computed by dividing the biopsy area by the total number of sequenced melanocytes. Consequently, the circle’s size is proportionate to the area covered by the enclosed cells. Due to geometric disparities between squares and circles, cases with multiple subclones might result in circles extending beyond the square boundaries. In such scenarios, the sizes of all circles within the square were proportionally reduced to fit within the square outline. Melanocytes carrying pathogenic mutations were marked in red, along with the outline of their enclosing circle. In instances where multiple cells shared somatic mutations, the largest circle represented the trunk of the phylogenetic tree with the subsequent circles within depicting the branches.

### Mixed-effects modeling

Mutation burden was compared between the tanning cohort and the combined data from the two control cohorts. Inference was performed at the cell level using a mixed model. That mixed model had a fixed effect for cohort (tanning versus non-tanning) and anatomic site (lower back versus upper back) and a random effect for subject. The likelihood ratio test was utilized to test the null hypothesis that the coefficient for cohort was equal to zero. Statistical significance corresponded to a *p*-value<0.05.

### Estimation of body surface area (related to figures 1C and S5)

The body surface area of various anatomical sites was approximated using the Lund and Browder chart^25^. For epidemiological investigations (Figure 1C and S5), we categorized the face/head/neck, arms below the elbows, and lower extremities below the knees as high CSD regions, as they are more likely to remain uncovered and thus are chronically exposed to sunlight. Conversely, regions typically shielded by clothing, such as the upper arms, front and back of the torso, and thighs, were defined as low CSD sites. The total body surface area for high and low CSD regions was calculated by summing the anatomical sites mentioned above.

