## Supplementary material for "Molecular effects of indoor tanning": Figure S1

### Fig. S1

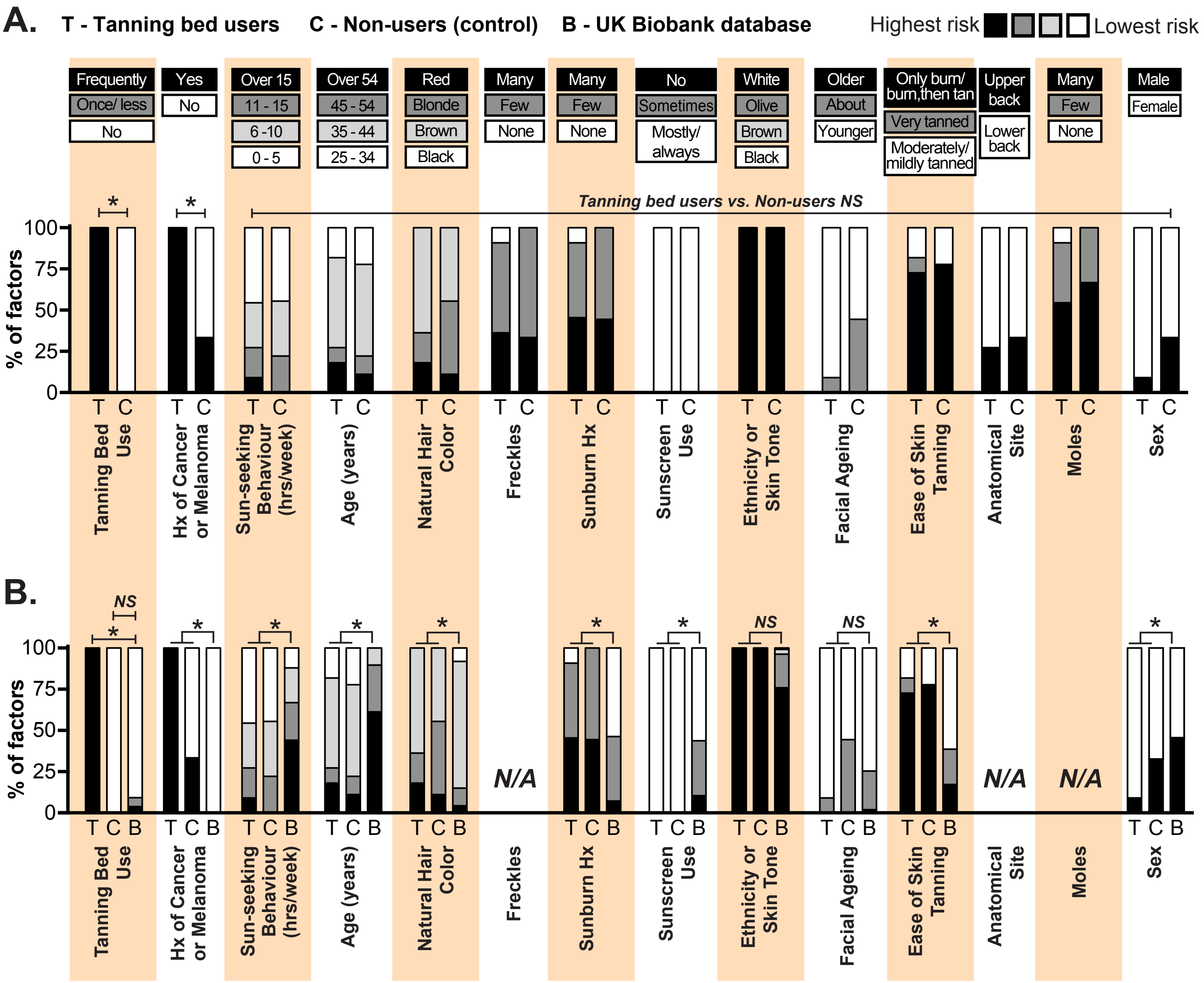


**Figure S1. Control cohort 1 donors match the risk profiles of tanning bed donors, however, both groups are at higher risk of skin cancer than the general population.** Donors were recruited for molecular studies of their skin cells and asked to fill out questions, modeled after the UK Biobank, surveying their past exposure to UV radiation and other risk factors for skin cancer. See Table S1 for the full wording of each question and answer. Bar graphs summarize the responses to each question with the risk trait colored more darkly. Asterisk denotes *p*-values less than 0.05 (Chi-squared test). Panel **A** compares the responses of the tanning cohort to the control cohort. Panel **B** compares the responses of both the tanning/control cohorts to the UK Biobank participants, chosen to represent the general population.
