## Supplementary material for "Molecular effects of indoor tanning": Figure S2

### Fig. S2

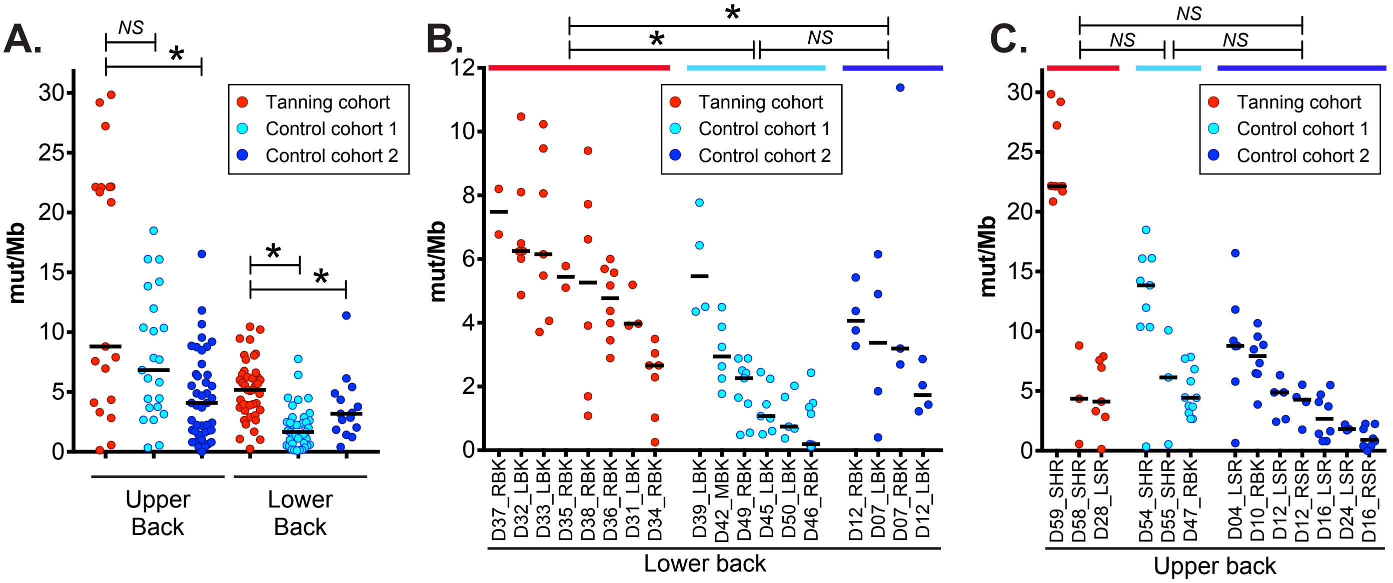


**Figure S2. Comparison of mutation burdens of melanocytes from tanning bed users to each control cohort separately.** Each data point corresponds to the mutation burden (measured in mutations per megabase) of an individual melanocyte. Black bars indicate median mutation burdens. Asterisks denote *p*-values less than 0.05 and *NS* indicate not significant (Wilcoxon rank-sum test). **A.** A comparison of melanocytes from tanning bed donors to melanocytes from each control cohort, separately for each anatomic site. **B.** A comparison of biopsy mutation burdens from tanning bed donors to control donors from each cohort on the lower back. The mutation burden of each biopsy was calculated from the median mutation burden of its constituent melanocytes. **C.** A comparison of biopsy mutation burdens from tanning bed donors to control donors from each cohort on the upper back.
