## Supplementary material for "Molecular effects of indoor tanning": Figure S3

### Fig. S3

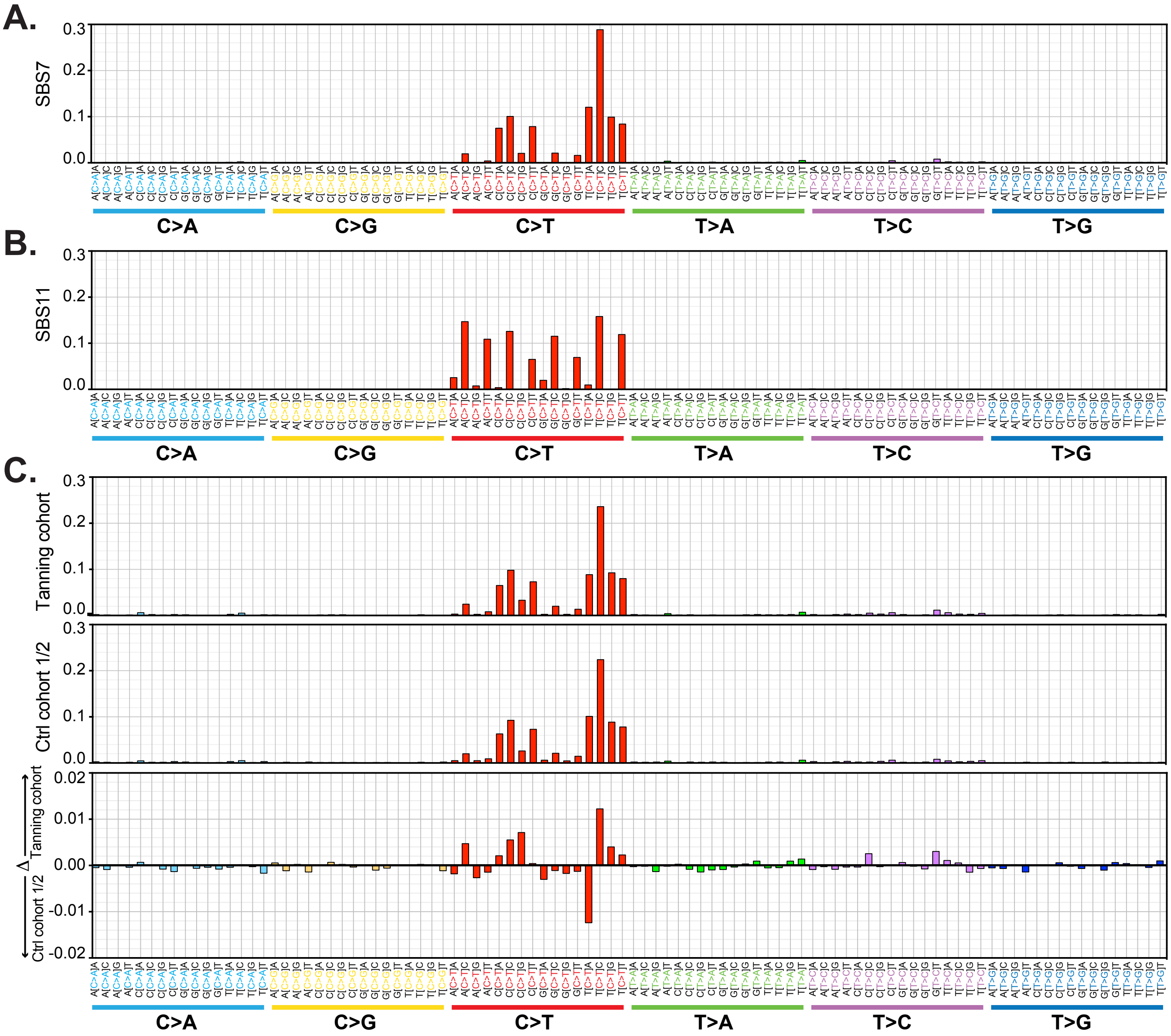


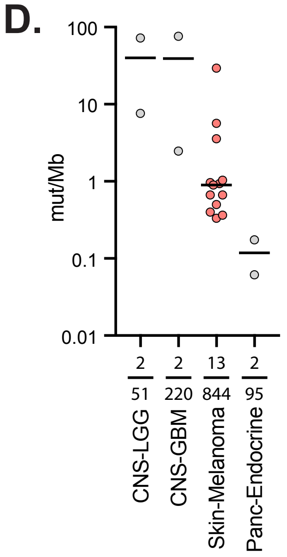


**Figure S3. Composition of mutational signatures.** 96 bar plots show the fraction of different mutations (y-axis) in all possible tri-nucleotide contexts (x-axis) for signature 7 (panel **A**) and signature 11 (panel **B**). Panel **C** compares the trinucleotide contexts of the tanning cohort (top panel), control cohort 1/2 (middle panel), and the difference between these two cohorts (bottom panel). Panel **D** illustrates the mutational burden attributable to signature 11 in the current catalog of COSMIC signatures (version 3.4).
