## Supplementary material for "Molecular effects of indoor tanning": Figure S4

### Fig. S4

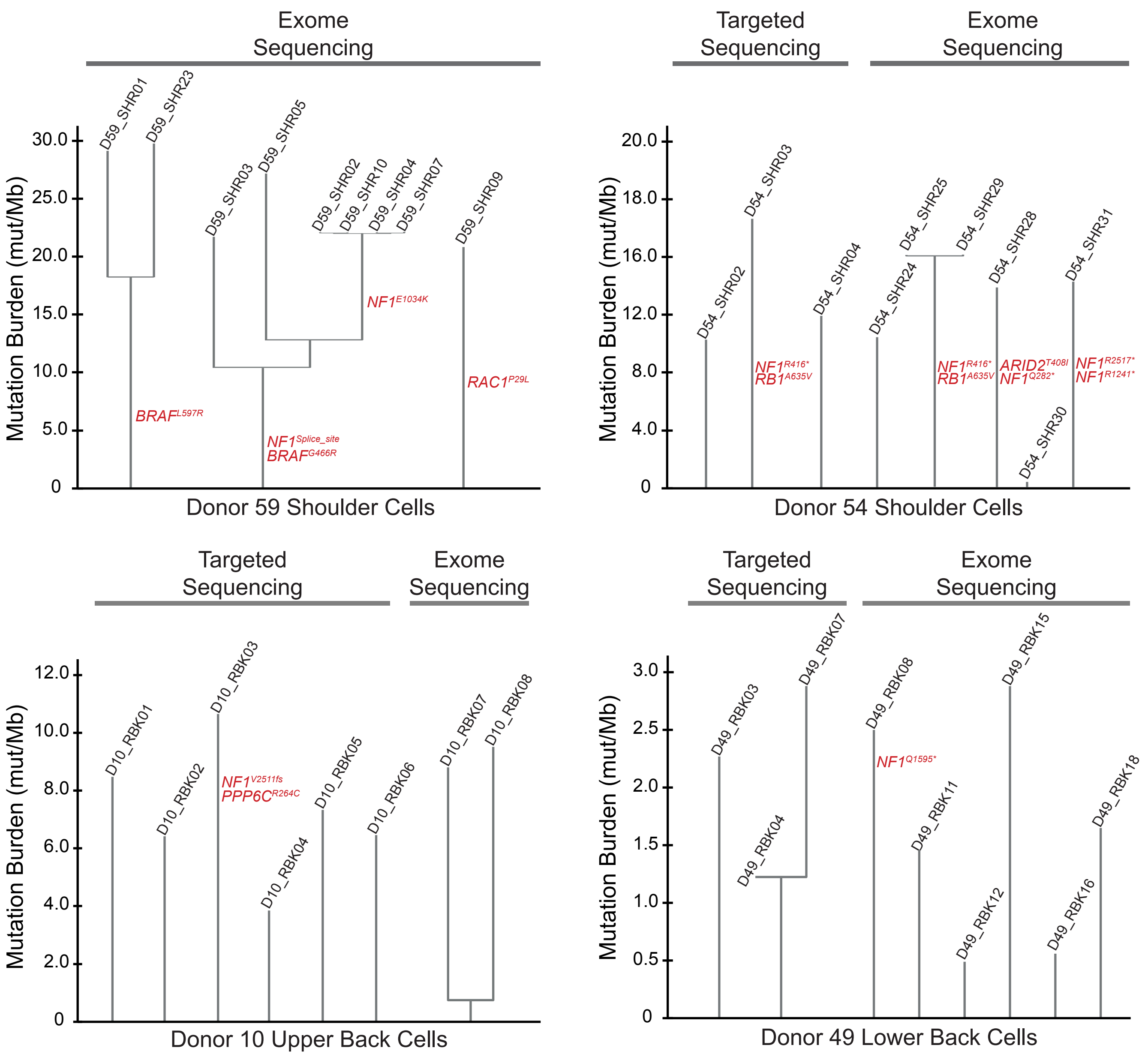


**Figure S4. Phylogenies of melanocytes from four skin biopsies with clonally related cells.** Phylogenetic trees were constructed for individual melanocytes from four skin biopsies. Each tree is rooted in the germline state. Trunk and branch distances are proportional to the number of shared and unshared somatic mutations between cells. The identity of each cell is labeled at the terminus of each tree, and pathogenic mutations are labeled in red. Some cells were sequenced with exome baits, and some were sequenced with a smaller gene panel, as indicated. See figure 4D for a series of schematics portraying the potential clonal structure of each biopsy.
