## Supplementary material for "Molecular effects of indoor tanning": Figure S5

### Fig. S5

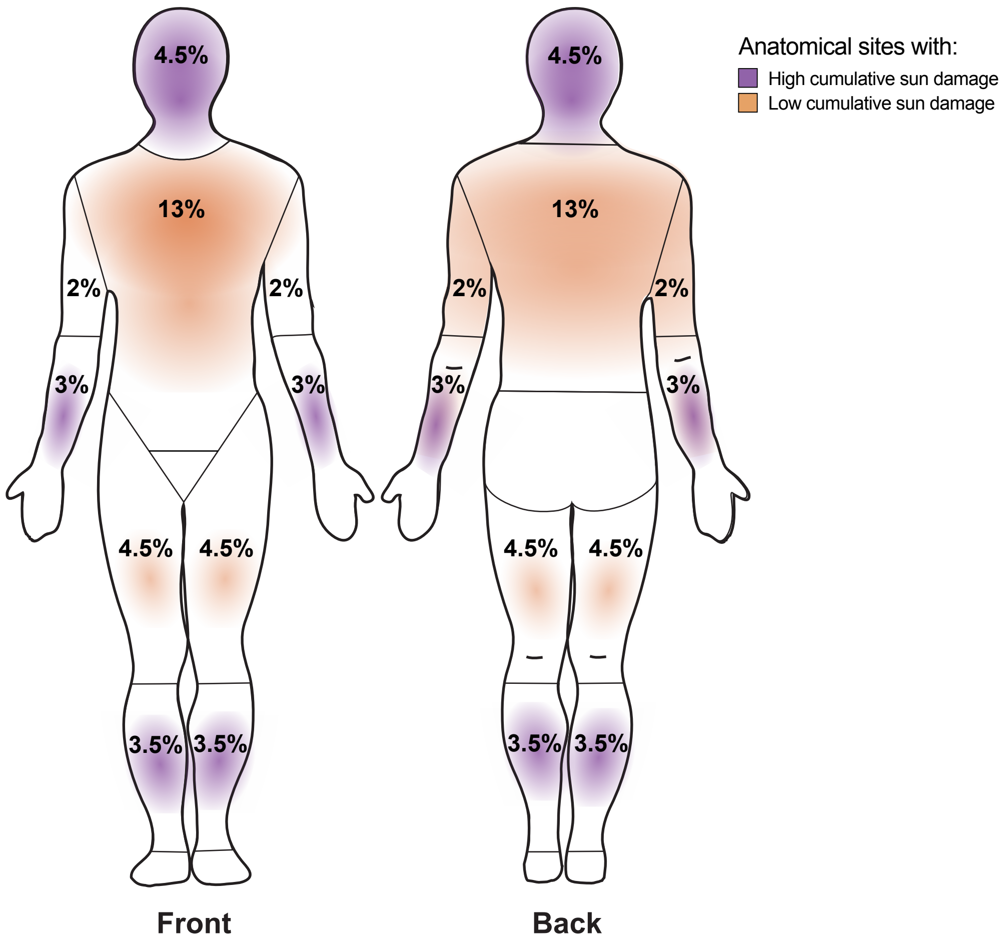


**Figure S5. Total body surface area in the regions with high (purple) and low (orange) cumulative sun damage (CSD).** The figure is based on Lund and Browder chart. The high CSD regions cover 35% and low CSD regions cover 52% of the body area. The remaining anatomical regions are considered sun shielded.
